# HDAC6-Mediated FoxO1 Acetylation And Phosphorylation Control Periodontal Inflammatory Responses

**DOI:** 10.1101/2024.12.10.627820

**Authors:** Hannah Lohner, Xiao Han, Junling Ren, Shuang Liang, Ruqiang Liang, Huizhi Wang

## Abstract

Post-translational modifications (PTMs) are critical regulators of protein function and cellular signaling. While histone deacetylation by histone deacetylases (HDACs) is well established, the role of specific HDACs in modulating non-histone protein PTMs, particularly in an infectious context, is poorly understood. Here, we reveal a pivotal role for HDAC6 in orchestrating periodontal inflammation through its dual regulatory effects on FoxO1 acetylation and phosphorylation. Using *Porphyromonas gingivalis*, a key periodontal pathogen, as a model pathogen, we observed that infection induces HDAC6 activation, driving inflammatory responses via modulating FoxO1 activity. HDAC6 depletion increased FoxO1 acetylation and phosphorylation, leading to its cytoplasmic sequestration and subsequent suppression of FoxO1- mediated pro-inflammatory cytokine production in macrophages. Mechanistically, HDAC6 deficiency not only directly enhances the acetylation of FoxO1 but also upregulates the expression of Rictor, a critical component of the mTORC2 complex, thereby promoting Akt phosphorylation and subsequently FoxO1 phosphorylation. This results in its cytoplasmic retention and attenuated inflammatory transcriptional activity. Functional studies demonstrated that HDAC6 depletion suppressed the production of key inflammatory mediators, including TNFα, IL-6, IL-12p40, and MIP-2, while promoting macrophage polarization toward anti-inflammatory M2 phenotypes. *In vivo*, using oral gavage infection and ligature-induced mouse periodontitis models, HDAC6 deficiency significantly reduced inflammatory cell infiltration in gingival tissues and protected against alveolar bone loss. These findings establish HDAC6 as a central regulator of periodontal inflammation, acting through the coordinated modulation of FoxO1 acetylation and phosphorylation. Beyond its role in oral pathology, HDAC6 may serve as a promising therapeutic target for managing inflammatory diseases linked to immune dysregulation.

## Introduction

Acetylation and deacetylation of histone proteins are dynamic and reversible processes that play critical roles in regulating inflammatory gene transcription by altering chromatin architecture (1–3). Deacetylation is mediated by histone deacetylases (HDACs), a family of enzymes that catalyze the removal of acetyl groups from lysine residues, thereby influencing protein structure, function, and interactions. HDACs are categorized into four classes: class I (HDAC 1, 2, 3, 8), class II (HDAC 4, 5, 6, 7, 9, 10), class III (Sirt1–Sirt7), and class IV (HDAC 11) (4, 5). The deacetylation of histone proteins exposes positively charged lysine residues, tightening chromatin structure and repressing transcriptional activity. Beyond histones, a growing body of evidence highlights the acetylation of non-histone proteins, including transcription factors involved in inflammation (6). Acetylation of these proteins at lysine residues often modulates their charge, conformation, and DNA-binding ability (7, 8). For example, deacetylation can enhance transcriptional activity by tightening interactions with DNA, which is negatively charged. Recent advances in HDAC inhibitors have underscored their therapeutic potential. A notable example is Givinostat (DUVYZAT™), a pan-HDAC inhibitor approved by the U.S. FDA in March 2024 for treating Duchenne Muscular Dystrophy. In addition to its efficacy in this context, Givinostat and other pan-HDAC inhibitors have demonstrated anti-inflammatory effects in various *in vitro* and *in vivo* models (9–14). These inhibitors underscore the need for a deeper understanding of the specific roles of individual HDACs in diverse physiological and pathological processes to inform the development of more targeted and effective therapeutics.

Forkhead box protein O1 (FoxO1), a member of the forkhead transcription factor family, is characterized by a conserved forkhead DNA-binding domain. While its precise functions remain incompletely understood, emerging evidence highlights FoxO1 as a central regulator of inflammation(15–17). FoxO1 drives the production of diverse pro-inflammatory mediators, including cytokines, proteolytic enzymes, and extracellular matrix regulators (18). Our and others studies, have shown that the nuclear translocation of FoxO1 is essential for its inflammatory activity in innate immune cells (16, 19, 20). FoxO1 phosphorylation at specific residues modulates its subcellular localization and activity; for instance, phosphorylation at serine 256 promotes its cytoplasmic retention, thereby attenuating its activity (16, 19–21) . In addition to phosphorylation, acetylation of FoxO1 at lysine residues significantly impacts its function. Acetylation promotes its cytoplasmic retention, preventing access to nuclear DNA and impairing its transcriptional activity (15, 22, 23). Conversely, FoxO1 deacetylation enhances its nuclear localization and activity, facilitating the expression of its target genes, as demonstrated in liver cells (24, 25). Considering the different post-translational modifications (PTMs), mediated by diverse enzymatic pathways, can exert convergent effects on FoxO1 activity, it is plausible that interplay or synergy between phosphorylation and acetylation exists. However, the regulation of FoxO1 acetylation and phosphorylation in periodontal inflammation, and their potential interactions in response to the challenge of oral pathogen, such as *P. gingivalis,* remains entirely unexplored.

Periodontitis is a polymicrobial infection-induced chronic inflammatory disease affecting over 38% of adults aged ≥30 years and nearly 90% of those aged ≥65 years (26). Emerging evidence indicates that periodontitis not only causes local tissue destruction but also increases the risk of systemic conditions, including atherosclerosis, diabetes, rheumatoid arthritis, Alzheimer’s disease, and oral cancers (27–30). The colonization of oral pathogens disrupts the homeostasis of the oral microbiome, triggering excessive inflammatory responses, leading to irreversible gingival tissue damage, alveolar bone loss, and the progression of periodontitis. Pathogen-associated molecular patterns (PAMPs) from oral bacteria activate immune receptors, driving the production of pro-inflammatory mediators such as TNF-α, IL-12, IL-1β, IL-6, and IL-8, which perpetuate periodontal inflammation (31–35). Consistent with the understanding that tissue destruction in periodontitis arises from excessive inflammation, antagonists targeting IL-1 and TNF-α have been shown to mitigate the severity of experimental periodontitis (36–38). These findings suggest the critical role of inflammation in periodontal tissue destruction and suggest that novel therapeutic strategies aimed at attenuating inflammatory responses to oral microbiota may offer effective approaches not only for controlling periodontitis but also for managing other chronic inflammatory diseases. Among the most well-studied oral pathogens, *Porphyromonas gingivalis* (*P. gingivalis*, Pg) is a keystone pathogen in periodontitis, contributing to the dysbiosis of the oral microbiota and the dysregulation of inflammatory immune responses. *P. gingivalis* employs a repertoire of virulence factors, including lipopolysaccharide (LPS), fimbriae, protease enzymes, and peptidoglycan proteins, to induce host inflammatory responses (39). Consequently, it is frequently used as a model organism for generating inflammation models that mimic the infectious microenvironment of periodontitis. Using a *P. gingivalis*-mediated infection model, we have previously demonstrated that inhibition of glycogen synthase kinase 3 (GSK3) (40) or overexpression of serum and glucocorticoid-induced kinase (SGK)-1 (16, 41) significantly reduces pro-inflammatory cytokine production and protects against periodontal bone loss. These findings suggest that identifying additional endogenous inflammatory regulators could pave the way for novel therapeutic strategies to mitigate the inflammatory responses. In this context, a pioneering study utilizing a pan-HDAC inhibitor demonstrated suppression of osteoclast development and a concomitant reduction in periodontal bone loss in a mouse model (42). Additionally, altered abundance of specific HDAC members has been observed in gingival tissues from patients with periodontitis (43). However, the immunoregulatory functions of individual HDACs in periodontal inflammation remain less known, and the underlying molecular mechanisms involved are yet to be elucidated. Given the controversial regulatory effects of HDACs on Toll-like receptor (TLR)- mediated pro-inflammatory gene expression (2, 44–46), individual HDAC members may exhibit distinct roles in controlling periodontal inflammation.

Unlike other HDACs, HDAC6 and HDAC10, which are predominantly localized in the cytoplasm, primarily deacetylate non-histone proteins and integrate multiple immune signaling pathways(47, 48). HDAC6 is particularly intriguing due to its capacity to regulate the activity of non-histone proteins, including transcription factors, through post-translational modifications (49–51). Deacetylated FoxO1 has been shown to enhance its transcriptional activity by facilitating nuclear localization, whereas its acetylation promotes cytoplasmic retention and impairs DNA-binding ability (15). Despite its established role in other inflammatory contexts, the regulation of FoxO1 acetylation and phosphorylation in periodontal inflammation, particularly in response to *P. gingivalis* infection, remains poorly understood. Furthermore, recent studies have shown that HDAC6-selective inhibitors can reduce the production of pro-inflammatory cytokines such as TNFα, IL-6, and IL-1β in sepsis models with minimal cytotoxicity (52–55), underscoring the therapeutic potential of targeting HDAC6 in inflammatory diseases. However, HDAC6’s specific activity and downstream signaling pathways in the context of oral bacterial infection and periodontal inflammation remain unexplored.

In this study, we employed cultured cell models along with two distinct periodontitis animal models to investigate the regulatory role of HDAC6 in periodontal inflammation. Our findings revealed a dual regulatory strategy employed by HDAC6, involving both direct acetylation and indirect phosphorylation mechanisms, to modulate oral bacteria-induced periodontal inflammation. Additionally, we mapped the signaling network downstream of HDAC6, elucidating its role in controlling FoxO1 phosphorylation and subsequent cytoplasmic sequestration during *P. gingivalis*-induced periodontal inflammation.

## Material and Methods

### Mice, bacteria, and reagents

10- to 12-week-old C57/BL/6J wild type (#000664) and *Hdac6^em2Lutzy^/J* (HDAC6 KO, #029318) male and female mice were purchased from the Jackson Laboratory and were maintained according to the animal care and standards of the Virginia Commonwealth University (VCU) animal facilities. All experiments were approved by the Institutional Animal Care and Use Committee (IACUC) at VCU (Protocol number, AD10002142) and were performed in strict accordance with standard operating procedures (SOPs) related to animal ethics and welfare. All efforts were made to minimize the number of mice used and to prevent animal distress, pain, and injury. Carbon dioxide (CO2) was used for euthanasia of mice. *Porphyromonas gingivalis* 33277 was from ATCC and cultured anaerobically in trypticase soy broth supplemented with yeast extract (1 mg/ml), hemin (5 μg/ml) and menadione (1 μg/ml). *Streptococcus gordonii* DL1 (from ATCC) were grown at 37°C in Anoxomat jars (Spiral Biotech) under microaerobic conditions (7% H2, 7% CO2, 80% N2, and 6% O2) in brain-heart infusion (BHI; Bacto, Sparks, MD) broth. Ultrapure LPS from *E. coli 0111:B4* and Pam3CSK4 were from Invivogen (San Diego, CA). Phospho-Serine antibodies were from Santa Cruz Biotechnology (Santa Cruz, CA). Total HDAC6 antibodies were from Abcam (Cambridge, MA). All other antibodies were from Cell Signaling Technology (Danvers, MA). The FoxO1 inhibitor AS1842856 was from MedChem Express (Monmouth Junction, NJ). The proteasome inhibitor MG-132 (also known as carbobenzoxyl-L-leucyl-L-leucyl-L-leucinal; ZLLL-CHO) was from SelleckChem (Houston, TX). Non-targeting pools of siRNA and a mixture of four pre-validated siRNA duplexes specific for *hdac6* (ON TARGET- *plus*^TM^) were from GE-Healthcare Dharmacon (Pittsburgh, PA). Mouse TNFα, IL-6, and IL-12P40 cytokine ELISA kits were purchased from Biolegend (San Diego, CA). Macrophage inflammatory protein 2 (MIP2)/CXCL2 kit was from R&D systems (Minneapolis, MN). RNeasy Mini and RNase-free DNase Set were from QIAGEN (Hilden, Germany). High-Capacity cDNA reverse transcription kit and Taq-path qPCR master mix were from Applied Biosystem (Foster City, CA).

### BMDMs and Mouse Embryo Fibroblasts (MEFs)

Bone marrow-derived macrophages (BMDMs) were generated from femoral and tibial bone marrow cells as previously described (56). Briefly, bone marrow was flushed from the femur and tibiae of 10 to 12-week-old *hdac6^-/-^* or the littermate control mice using sterile Hanks’ balanced salt solution and homogenized by passage through an 18.5-gauge syringe repeatedly. The cells were washed in PBS, centrifuged at 1500 rpm for 5 min, and resuspended in RPMI 1640 medium supplemented with 10% FBS, 50 μM 2-mercaptoethanol, 1 mM sodium pyruvate, 2 mM L-glutamine, 20 mM HEPES, 50 units/ml penicillin, 50 μg/ml streptomycin, and 30% L929 culture supernatant. Nonadherent cells were collected after 24 hours and cultured for 7 days in Costar ultra-low attachment polystyrene culture dishes with a medium change on day 4. Macrophages were about 95% F4/80^+^/CD11b^+^ as determined by flow cytometry and ready for further experiments. Mice primary oral epithelial cell were purchased from Lifeline Cell Technology (Frederick, MD), and maintained in our lab according to the manufacturer’s procedure. Mouse embryonic fibroblasts (MEFs) (ATCC, #SCRC-1008) were cultured in DMEM (ATCC, #30-2002) modified with 4mM L-glutamine, 4500mg/L glucose, 1mM sodium pyruvate, 1500mg/L sodium bicarbonate, 15% FBS, and 50 units/mL of penicillin and streptomycin. Medium was changed twice a week or when the pH changed. Rictor deficient MEFs were obtained from Dr. Hongbo Chi’s lab at St. Jude Children’s Research Hospital (Memphis, TN).

### siRNA transfection, cytokine assay, and Western blots

siRNA and plasmid transfections in human monocytes were carried out by electroporation using a Nucleofector device (Amaxa, Germany) according to the proposed protocol. Briefly, 4 × 10^6^ MEFs were re-suspended in 100 μl of Nucleofector solution (MEF Nucleofector kit; Amaxa) together with 2μg of a green fluorescent protein (GFP)-coding plasmid (pCMV-GFP) and 2μg of siRNA duplexes for each target. After electroporation, 400μl of pre-warmed M-199 containing 10% FCS was added to the cuvette and the cells were transferred into culture plates containing pre-warmed M-199 with 10% FCS. At the optimal time of gene silencing (48 hours post-transfection), cells were exposed to *P. gingivalis*. BMDMs were transfected with nontargeting control siRNA and siRNA-*hdac6* using Lipofectamine RNAiMAX (Invitrogen, Carlsbad, CA) following the manufacturer’s protocol. After transfection, the cells were directly seeded in either 96- or 6-well plates. The levels of HDAC6 were assessed by Western blots. Cells were lysed in RIPA buffer containing phosphatase inhibitors for Western blot assays. Images were acquired using the G:Box Chemi XXI (Syngene, Cambridge, UK). For cytokine assays, cell-free supernatants were collected after stimulation with *P. gingivalis* 12 hours instead of 24 hours to minimize gingipain-induced proteolytic cleavage of cytokines. Cytokine concentration was determined by enzyme-linked immunosorbent assay (ELISA) following the manufacturer’s instructions (Biolegend, San Diego, CA).

### RNA isolation, real-time quantitative PCR (qRT-PCR), cell fraction, and immunoprecipitation

Total RNA was isolated with the RNeasy Mini Kit (Qiagen, #74106) and DNase Set (Qiagen, #79256) as per the manufacturer’s guidelines. Samples were reverse transcribed using the High-Capacity cDNA reverse transcription kit (Thermofisher, # 4368813). Real-time quantitative PCR analysis was performed in triplicate. PCR was performed in an Applied Biosystems 7500 system using SYBR Green master mix (Thermofisher, #4367659). Relative levels of gene expression were determined using GAPDH and β-actin as controls and measured using the ΔΔCT method. For the cell fractionation preparation, 4.7 x 10^6^ BMDMs were seeded into 60mm dishes and allowed to become adherent (overnight). Cytoplasmic and nuclear cell fractions were collected using the NE-PER Nuclear and Cytoplasmic Extraction kit (Thermofisher, #78833). Western blot analysis was then implemented as described above. GAPDH and Lamin B1 were used as loading controls for the cytoplasmic and nuclear fractions respectively. For immunoprecipitation, 10-20 x 10^6^ BMDMs were seeded into 15 cm dishes and allowed to become adherent (overnight). Samples were processed using a Pierce Classic Magnetic IP/Co-IP Kit (Thermofisher, #88804) as specified by the manufacturers guide. Briefly, cells were treated with *P. gingivalis* for the appropriate timepoints, supernatant was removed, and lysis buffer containing ReadyShield Phosphatase and Protease inhibitor cocktail was added to the cells. Samples were kept on ice for 5 minutes and then centrifuged for 10 minutes at 13,000 *× g* at 4°C. Protein concentration of the supernatants was measured with the BCA Assay. Cell lysates up to 1000µg of protein were combined with HDAC6 antibody and turned overnight at 4°C to form an immune complex. The immune complex was combined with 25µL of Pierce Protein A/G Magnetic Beads (Thermofisher, #88802) and turned at room temperature for 1 hour. The beads were collected with a magnetic stand and bound samples were eluted using the alternative elution method per the protocol instructions. Eluted samples were run for Western blot analysis.

### *P. gingivalis*-induced periodontal inflammation model and immunohistochemistry

The endogenous oral microbiota was suppressed in 10- to 12-week-old wild type and HDAC6 knockout mice by sulfamethoxazole (800 μg/ml) and trimethoprim (400 μg/ml) provided *ad libitum* in water for 5 days. The mice then received pure drinking water for 5 days. Alveolar bone loss was induced by oral infection with 1 × 10^9^ CFU of *P. gingivalis* suspended in 100 μl of phosphate-buffered saline with 2% carboxymethylcellulose. Infections were performed six times at every other day. Wild type and HDAC6 knockout mice were euthanized with CO2 and cervical dislocation 42 days after the final infection. Maxillary gingiva from mice upper jaws were harvested with a half used for RT-PCR assay, and the other half for Western blot assay. The fresh gingival tissues were immersed in RNAlater® (Ambion Cat. AM7020) or RIPA buffer with protease and phosphatase inhibitors (Millipore Sigma, Cat. P8340 and P0044) (1:100) and then stored at -20°C for further RT-PCR or Western blot assay, respectively. The lower jaws of the mice were fixed in 4% paraformaldehyde, decalcified in Immunocal solution for 15 days and embedded in paraffin wax for immunofluorescence assays. Alveolar bone loss was measured via microcomputed tomography scanning (as shown below). The results were expressed as the mean with S.D.. The paraffin embedded tissue blocks were freshly cut into 4 μm mesiodistal sections for subsequent immunostaining with hematoxylin and eosin (H&E) to evaluate inflammatory cell infiltration, and Alexa 488-conjugated anti-mouse Ly6G was used to measure the infiltration of neutrophils.

### Immunofluorescence (IF) for Cells

2 x 10^5^ BMDMs were seeded onto a round cover slip seated inside the well of a 24-well plate and were allowed to adhere to the cover slip (overnight). The BMDMs were challenged with WT Pg and collected from 30 minutes to 24 hours post infection. Cells were fixed in 4% paraformaldehyde solution in PBS (PFA) (Santa Cruz, #SC-281692) for 15 minutes and permeabilized in 0.5% Triton X-100 (Sigma Aldrich, #T8787) in PBS for 15 min at room temperature. Blocking was achieved with 3% BSA in PBS for 1 hour at room temperature. FoxO1 primary antibody in 3% BSA was incubated overnight at 4°C, followed by secondary Alexa Fluor 488 goat anti-rabbit in PBS for 1 hour at room temperature. F-actin was labeled with Alexa Fluor 549 Phalloidin in PBS for 1 hour at room temperature. The samples were mounted on microscope slides using ProLong Gold antifade reagent with DAPI (Thermofisher, #P36935), the edges were sealing with clear nail polish. The samples were visualized and images captured by a Keyence microscope.

### Ligature-Induced Periodontal Disease Model

Procedures were modified and performed based on previously described studies (57, 58). Briefly, 10-12-week-old WT and HDAC6 KO mice were anaesthetized with 2.5% isoflurane in oxygen. Mice were then transferred to a 3D-printed bed (Dr. Julie Marchesan, UNC) and stabilized with rubber bands. 5-0 suture silk (either soaked in a sterile PBS sham solution or solution of 1 x 10^9^ CFU *P. gingivalis*) was inserted on both sides of the left maxillary M2 molar and tied on the palatal side of the molar. After the procedure, the mice were allowed to recover on a controlled heat source until normal behavior returned. 3 days post ligature placement, mice received an oral inoculation of either 1 x 10^9^ CFU *P. gingivalis* in 2% CMC or CMC sham control. 14 days after ligature placement, mice were euthanized with CO2. Maxillae were fixed in 10% formalin solution for 48 hours and used to quantify bone loss via microcomputed tomography scanning (as shown below).

### Microcomputed tomography (micro-CT)

Mice from the oral infection induced- and ligature-induced periodontal disease models were euthanized at their appropriate endpoints. Mouse skulls were fixed in 10% formalin solution and used for microcomputed tomography (micro-CT) scanning on a Skyscan 1173 scanner (Micro Photonics Inc). The X-ray was run with a voltage of 92kV, current of 80µA, and exposure time of 1100ms. All reconstruction of scans was done through the NRecon software (Micro Photonics Inc.). Distance from the cemento-enamel junction to the alveolar bone crest (CEJ-ABC) was measured through the DataViewer software (Micro Photonics Inc.) from the sagittal plane. For the oral infection mice, the average of the measurements taken from 6 sections from the buccal and 6 sections from the palatal sides of the M1 molar were evaluated. For the ligature mice, the average of the measurements taken from 6 sections from the buccal and 6 sections from the palatal sides of the M2 molar were evaluated.

### Statistical analyses

The statistical significance of differences between groups was evaluated by the analysis of variance (ANOVA) and the Tukey multiple comparison test using the InStat program (GraphPad). Differences between groups were considered significant at the level of *P* ≤ 0.05.

### Data availability

All the raw data used for figure generation and any additional information are available upon request from Huizhi Wang.

## Results

### HDAC6 is Essential for the Production of Inflammatory Mediators in TLR Ligands- Stimulated Immune Cells

HDAC6 has been reported to play a dual role in regulating the production of inflammatory mediators (2, 44). Pan-HDAC inhibitors and siRNAs of the specific individual HDAC are widely used in previous studies (44, 52, 53). To avoid potential limitations associated with chemical inhibitors and siRNA, such as non-specific effects and off-target actions, we utilized bone marrow-derived macrophages (BMDMs) generated from wild-type (WT) and HDAC6 knockout (KO) mice to examine the functional role of HDAC6 in TLR activation induced inflammatory mediators. We first examined HDAC6 expression in macrophages and its deacetylase activity. As shown in Figure 1A, HDAC6 KO resulted in a complete loss of HDAC6 expression and a remarkable increase acetylation in α-tubulin, a specific substrate of HDAC6. These findings confirm the successful depletion of HDAC6 in KO mice and the specificity of α-tubulin acetylation as a marker of HDAC6 activity in BMDMs. To investigate the role of HDAC6 in inflammatory responses, we stimulated BMDMs with various Toll-like receptor (TLR) agonists, including *Escherichia coli* lipopolysaccharide (*E. coli* LPS) and Pam3CSK4 (Pam3A), for 24 hours. The expression levels of key inflammatory cytokines, IL-6, IL-12p40, interferon beta, and TNFα, were examined by ELISA or Western blot (Figure 1B, C). Our results showed that HDAC6 deficiency significantly reduced the production of these inflammatory mediators, indicating that HDAC6 is essential for their expression in TLR-stimulated immune cells.

**Figure 1.**
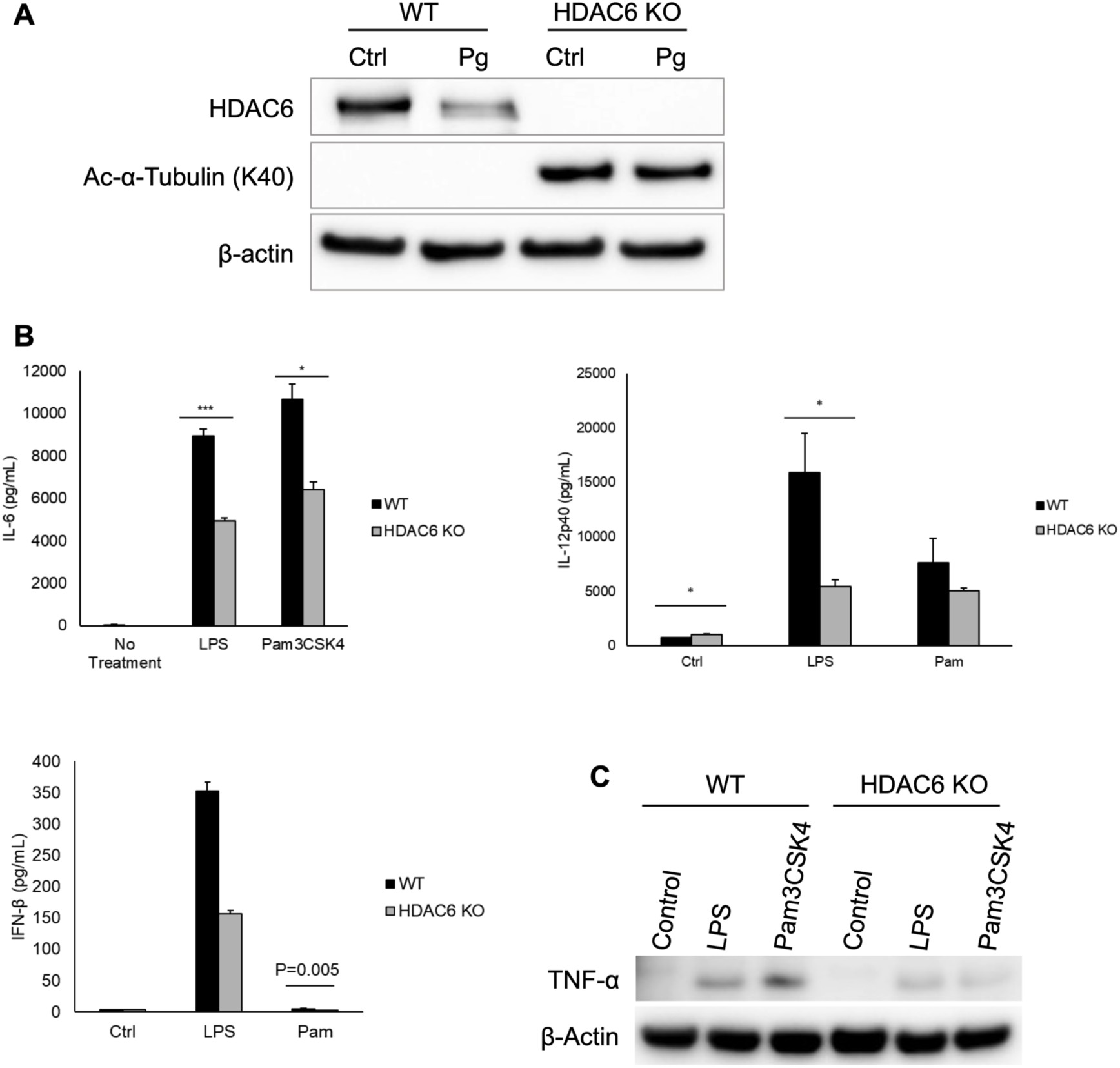
HDAC6 depletion attenuates TLR2- and TLR4-mediated cytokine protein secretion. **(A)** Western blot analysis confirms HDAC6 depletion in bone marrow-derived macrophages. HDAC6 expression is observed in wild-type (WT) cells with or without *P. gingivalis* infection, while HDAC6 knockout (KO) cells exhibit increased acetylation of α-tubulin, a known substrate of HDAC6; **(B)** ELISA quantification of cytokine secretion shows a significant reduction in IL-6, IL-12p40, and IFN-β levels in HDAC6 KO BMDMs following 24-hour stimulation with the TLR4 agonist LPS or the TLR2 agonist Pam3CSK4; **(C)** Western blot analysis of TNFα expression in whole-cell lysates of BMDMs 24 hours after TLR agonist treatment. *, and ***, represent the statistical significance with *P* < 0.05 and *P < 0.001,* respectively.

### HDAC6 is Activated in Response to *P. gingivalis* Challenge and Is Required for the Induction of Inflammatory Mediators

To investigate the functional role of HDAC6 in oral inflammation induced by infectious agents, we utilized *Porphyromonas gingivalis* (*P. gingivalis*) as a representative oral pathogen. Immunoprecipitation of HDAC6, followed by serine phosphorylation analysis, revealed that *P. gingivalis* stimulation induces serine phosphorylation of HDAC6 compared to untreated controls (Figure 2A). Intriguingly, pretreatment with the *P. gingivalis* gingipain inhibitor, TLCK, abolished serine phosphorylation of HDAC6, highlighting the essential role of gingipain in *P. gingivalis*- mediated HDAC6 phosphorylation. Notably, HDAC6 phosphorylation differentially affects its deacetylase activity (59–61). Phosphorylation of essential serine residues in the DD1/DD2 domain of HDAC6 such as Serine 22 or 458 is required for its activation, while phosphorylation of tyrosine residues such as Tyrosine 570 in other regions can negatively regulate deacetylase activity (62–65) (Figure 2B). Therefore, our results indicate *P. gingivalis* infection activates HDAC6 in macrophages. Additionally, a colorimetric assay demonstrated that *P. gingivalis* infection remarkably enhances HDAC6 enzymatic activity in BMDMs (Figure 2C). To examine the role of HDAC6 in the production of inflammatory mediators, we further quantified cytokine levels in *P. gingivalis*-stimulated WT and HDAC KO macrophages. HDAC6 depletion significantly reduced the production of IL-6 and TNFα at both message and protein levels (Figure 3A). Considering the critical role of chemokines, particularly CXCL2 (MIP-2), in neutrophil recruitment during inflammation, we also evaluated the production of MIP-2 in response to *P. gingivalis* challenge. Our results showed that HDAC6 depletion significantly decreased MIP-2 levels in *P. gingivalis*- stimulated BMDMs (Figure 3B). Collectively, these findings indicate that HDAC6 activation is required for the production of inflammatory cytokines and chemokines in response to *P. gingivalis*. This highlights HDAC6 as a critical regulator of periodontal inflammation.

**Figure 2.**
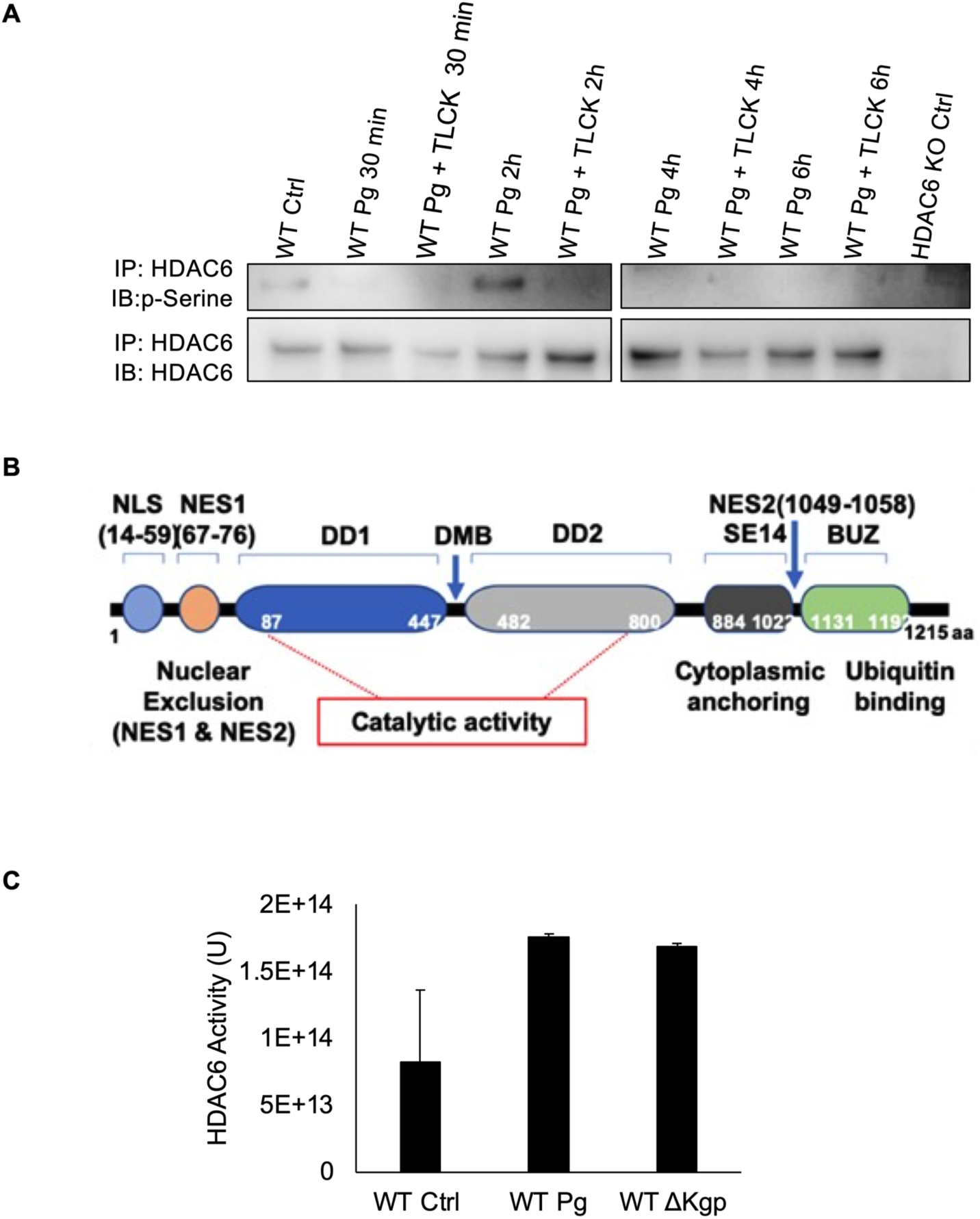
*P. gingivalis* infection enhances HDAC6 activity. **(A)** Co-immunoprecipitation (Co-IP) analysis of phosphorylated serine residues on HDAC6 in wild-type (WT) BMDMs infected with *P. gingivalis*. Cells were collected at the indicated time points post-infection, and samples from *P. gingivalis-*treated with the gingipain inhibitor TLCK were included for comparison. HDAC6 knockout (KO) BMDMs served as negative controls to validate HDAC6- specific phosphorylation. **(B)** Schematic model of HDAC6 domains; **(C)** Deacetylase activity of HDAC6 measured using an enzymatic assay kit (Abcam) 2 hours after WT BMDMs were challenged with either WT *P. gingivalis* or a gingipain deletion mutant.

**Figure. 3.**
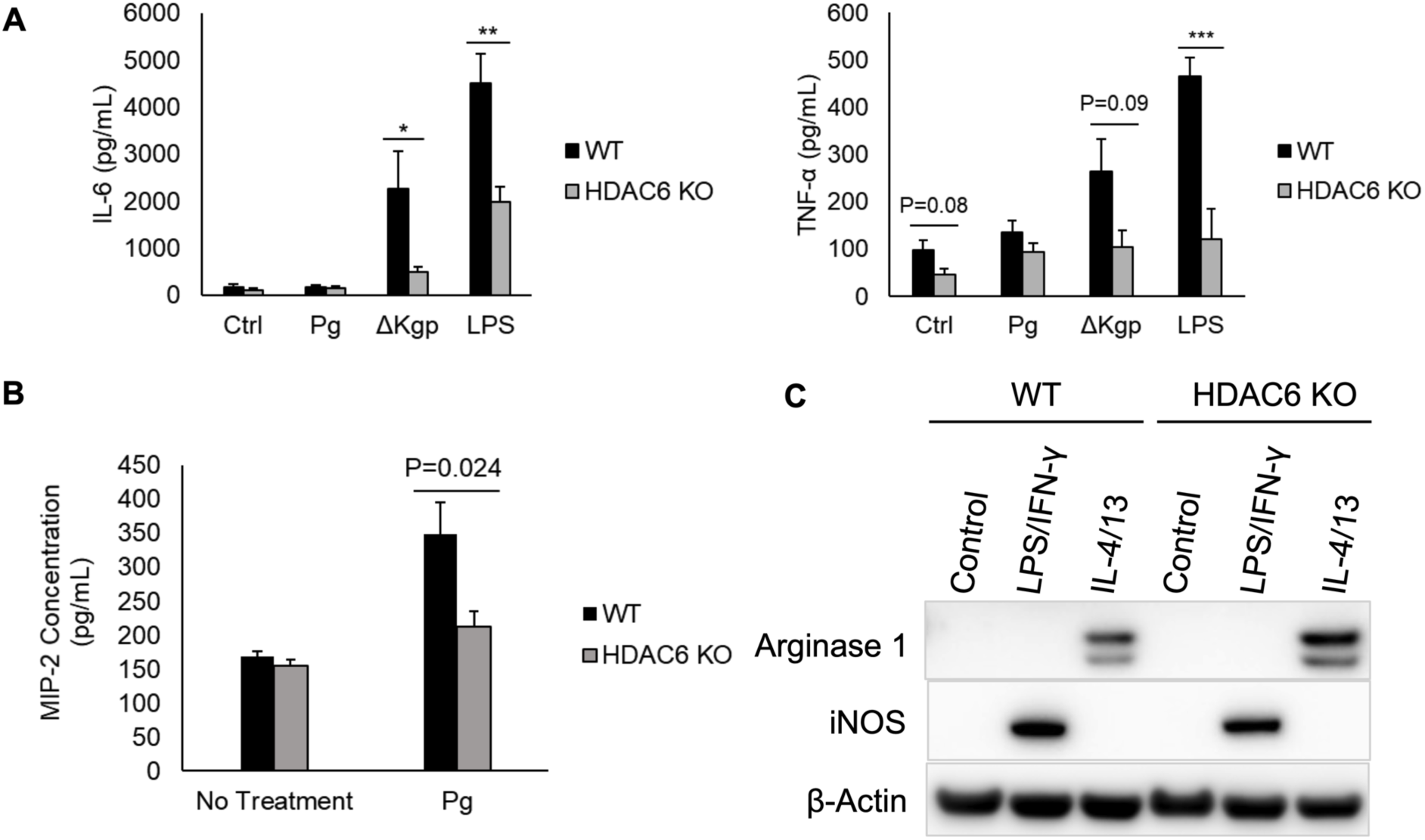
HDAC6 deletion reduces cytokine and chemokine secretion in response to *P. gingivalis* infection and skews macrophage polarization toward M2 phenotypes. **(A, B)** WT and HDAC6 KO BMDMs were challenged with WT *P. gingivalis*, a gingipain deletion mutant (ΔKgp), or *E. coli* LPS (positive control). ELISA assays were used to measure secreted protein levels of IL-6, TNF-α, and MIP-2 8 hours post-infection. **(C)** WT and HDAC6 KO BMDMs were treated with the indicated molecules for 24 hours. Western blot analysis was performed to detect M1- versus M2-associated markers (iNOS vs Arginase 1) using the indicated antibodies. *, **, and ***, represent the statistical significance with *P* < 0.05, *P<0.01,* and *P < 0.001,* respectively.

### HDAC6 Depletion Shifts Macrophage Polarization Towards M2 Phenotypes

Inflammatory responses are tightly regulated to maintain homeostasis through diverse mechanisms operating at multiple levels. Macrophage polarization is a pivotal event in the progression and resolution of inflammation. To examine the possible regulatory role of HDAC6 in macrophage polarization, we assessed characteristic marker expression in BMDMs derived from WT and HDAC6 KO mice. These macrophages were primed with *E. coli* LPS/IFN-γ to induce M1 polarization or IL-4/IL-13 to induce M2 polarization, following protocols established in our previous studies (16). As shown in Figure 3C, HDAC6 depletion significantly decreased the expression of inducible nitric oxide synthase (iNOS), a marker of M1 polarization, while concurrently increasing the expression of arginase 1 (Arg-1), a key marker of M2 polarization. iNOS catalyzes the production of nitric oxide (NO) from L-arginine, which is essential for driving macrophages towards an M1 phenotype characterized by pro-inflammatory responses. Conversely, Arg-1 converts L-arginine into urea and ornithine, promoting the anti-inflammatory M2 phenotype. Combined with the results in figure 1, showing HDAC6 depletion reduces the production of pro-inflammatory cytokines including IL-6, IL-12p40, and TNFα, these findings suggest that HDAC6 KO mitigates inflammatory responses by not only reducing the production of inflammatory cytokines but also by favoring macrophage polarization towards M2 phenotypes, thereby contributing to the resolution of inflammation.

### HDAC6 Reduces FoxO1 Acetylation and Enhances Its Nuclear Localization in Response to *P. gingivalis* Challenge

To further elucidate the downstream signaling pathways through which HDAC6 modulates inflammatory immune responses, we first examined the activation of prototypical inflammatory pathways, including NF-κB and MAPK. Our results demonstrated that HDAC6 depletion substantially reduces the activation of both pathways in *P. gingivalis*-stimulated macrophages at various time points (Figure 4A). As shown in figure 2 and 3, *P. gingivalis* kgp Δkgp mutant induces even higher activity of NF-kB and MAPK signaling pathways in both WT and KO macrophages, indicating this enzyme is crucial for *P. gingivalis* induced inflammatory responses. Unlike other HDACs, which are primarily located in nucleus, HDAC6 is predominantly localized in the cytoplasm and primarily targets non-histone proteins for deacetylation. We therefore investigated whether HDAC6-mediated deacetylation is involved in the regulation of *P. gingivalis*-induced inflammation.

**Figure 4.**
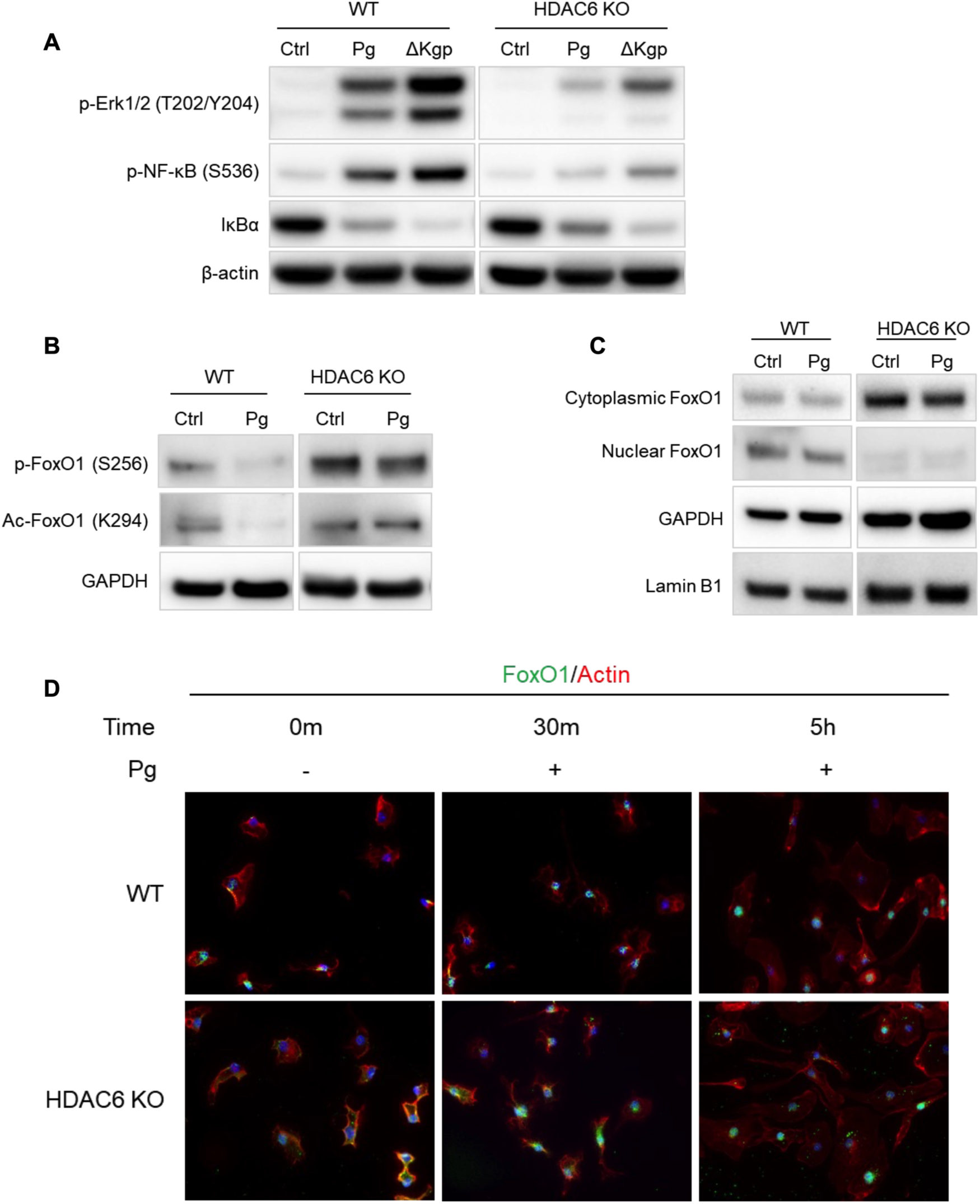
HDAC6 deletion reduces activation of downstream transcription factors and kinases while impairing nuclear shuttling of FoxO1. **(A)** WT and HDAC6 KO BMDMs were challenged with WT *P. gingivalis* or the gingipain deletion mutant (ΔKgp). Total cell lysates were collected 30 minutes post-infection and analyzed via Western blot for NF-κB and ERK activation using the indicated antibodies. **(B, C)** WT and HDAC6 KO BMDMs were infected with *P. gingivalis*. **(B)** Whole-cell lysates were collected 30 minutes post-infection to examine FoxO1 phosphorylation and acetylation. **(C)** Cytoplasmic and nuclear fractions were collected 24 hours post-infection and analyzed for FoxO1 localization and post-translational modifications by Western blot. **(D)** Immunofluorescence staining of BMDMs for FoxO1 (green), F-actin (red), and the nucleus (blue) at the indicated time points post-infection, visualized at 60× magnification.

FoxO1 is a transcription factor and known to regulate the production of inflammatory cytokines and macrophage polarization by shuttling between the cytoplasm and nucleus (16, 19, 20). Furthermore, FoxO1 has been shown to be deacetylated by other HDAC family members in liver cells (24, 25). To explore the possible role of HDAC6 in FoxO1 regulation, we assessed FoxO1 acetylation and subcellular localization in *P. gingivalis*-stimulated macrophages. As shown in Figure 4B, HDAC6 KO increased FoxO1 acetylation in unstimulated BMDMs, indicating that FoxO1 is a specific target of HDAC6. Furthermore, in WT BMDMs, *P. gingivalis* infection markedly reduced FoxO1 acetylation. while in HDAC6 KO BMDMs, *P. gingivalis* failed to do so (Figure 4B), strongly suggesting that HDAC6 is activated by *P. gingivalis*, which is essential for FoxO1 deacetylation. These findings also confirmed colorimetric results (Figure 2C) showing *P. gingivalis* enhances HDAC6 activity in macrophages.

We next examined FoxO1 subcellular localization and found that HDAC6 deficiency significantly reduced the nuclear abundance of FoxO1, resulting in its accumulation in the cytoplasm (Figure 4C). This observation was further validated through immunofluorescence staining (Figure 4D). Since nuclear FoxO1 is critical for its regulatory functions in inflammatory responses, these results suggest that HDAC6 deficiency promotes the cytoplasmic retention of FoxO1, thereby impairing its ability to regulate *P. gingivalis*-induced inflammatory responses in BMDMs.

### HDAC6 Suppresses Rictor Expression and Promotes FoxO1 Nuclear Localization by Limiting Akt Phosphorylation

The shuttling of FoxO1 between the cytoplasm and nucleus is primarily regulated by its phosphorylation at Ser256, a process mediated by upstream signaling pathways such as PI3K-Akt (19, 20). HDAC6 has been shown to bind with ubiquitin and thus implicate in protein ubiquitination processes (66, 67). To determine whether HDAC6 regulates the expression of signaling molecules involved in FoxO1 shuttling, we examined the expression of various components of the PI3K-Akt pathway. While most components showed no significant changes (data not shown), we observed a substantial increase in the expression of Rictor, an essential component of mTORC2 and a key regulator of Akt phosphorylation, in *P. gingivalis*-stimulated BMDMs lacking HDAC6 (Figure 5A). This finding was further validated using HDAC6-specific siRNA in mouse fibroblasts, which also enhanced Rictor expression (Figure 5B).

**Figure 5.**
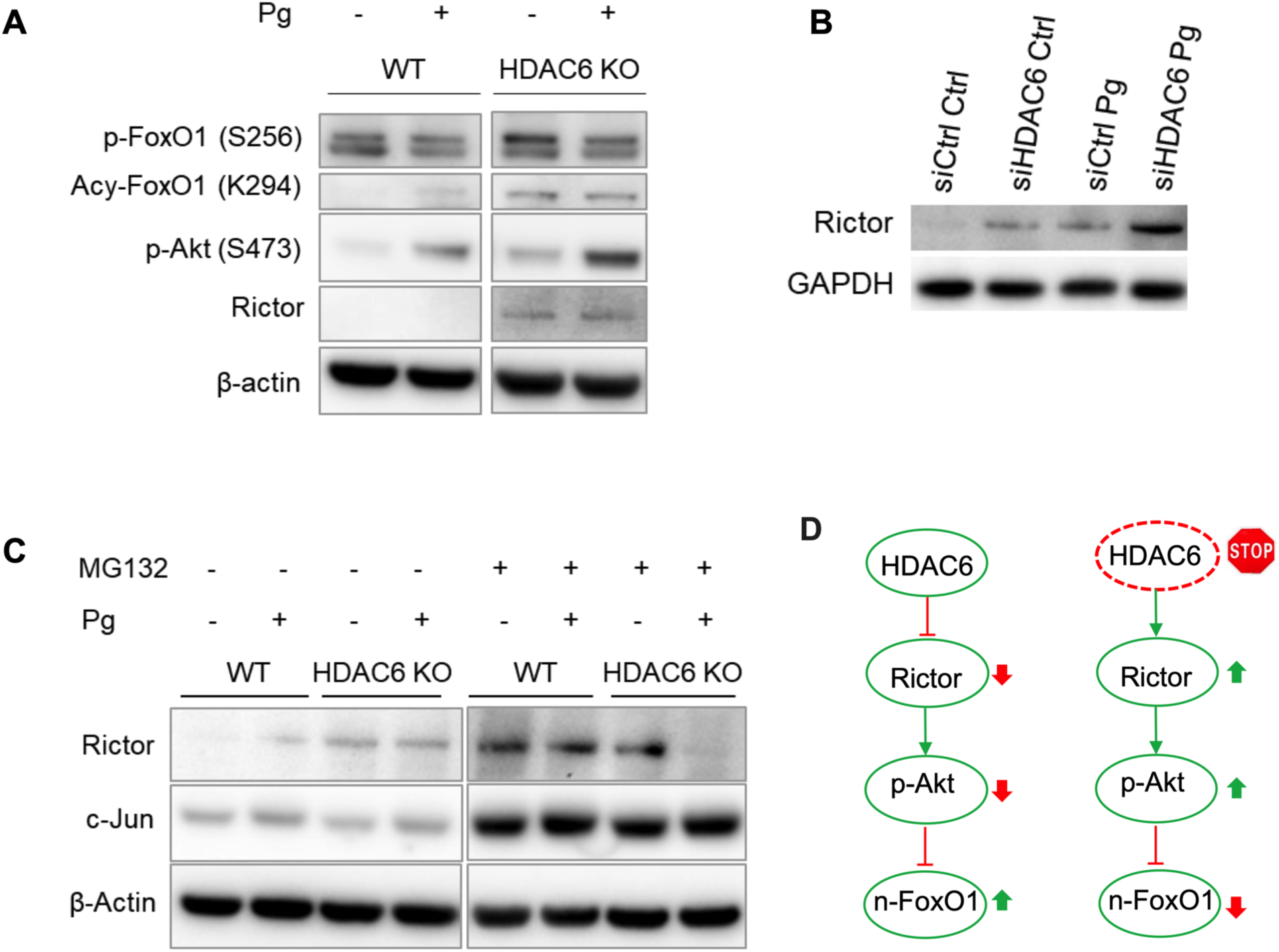
HDAC6 KO leads to a direct increase in FoxO1 acetylation and a Rictor-Akt-mediated increase in FoxO1 phosphorylation. **(A)** Whole-cell lysates from WT and HDAC6 KO BMDMs were collected 30 minutes post-*P. gingivalis* infection and analyzed by Western blot using the indicated antibodies. The data show that HDAC6 KO increases both the phosphorylation and acetylation of FoxO1. **(B)** Mouse fibroblasts treated with siRNA targeting *hdac6* were infected with *P. gingivalis*. Whole-cell lysates collected 30 minutes post-infection were analyzed by Western blot, demonstrating that HDAC6 silencing enhances Rictor expression in response to *P. gingivalis*. **(C)** WT and HDAC6 KO BMDMs were pretreated with the proteasome inhibitor MG132 for 1 hour prior to *P. gingivalis* infection. Whole-cell lysates were collected 30 minutes post-infection and analyzed by Western blot, showing that increased Rictor expression in HDAC6 KO cells is partially dependent on ubiquitination-mediated proteasome activity. Schematic model illustrating how HDAC6 knockout reduces nuclear FoxO1 abundance and activity by increasing both acetylation and Rictor-mediated phosphorylation, ultimately promoting FoxO1 cytoplasmic retention.

To explore the mechanism by which HDAC6 deficiency increases Rictor expression, we pretreated BMDMs with MG-132, a proteasome inhibitor (Figure 5C). MG-132 treatment substantially elevated Rictor expression in both wild-type and HDAC6 knockout macrophages, suggesting that HDAC6 deficiency enhances Rictor expression by limiting its ubiquitination and subsequent proteasomal degradation (Figure 5C). This observation is consistent with a recent study reporting that HDAC6 actively involve in the ubiquitination process to regulate the expression of specific proteins (68, 69) Moreover, HDAC6 deficiency-mediated increases in Rictor expression were consistently accompanied by elevated phosphorylation of Akt and FoxO1, as well as increased FoxO1 acetylation, in *P. gingivalis*-stimulated BMDMs (Figure 5A). Given that both phosphorylation of FoxO1 at Ser256 and its acetylation restrict FoxO1 to the cytoplasm (22, 23), these results suggest that HDAC6 deficiency suppresses FoxO1 activity by enhancing its phosphorylation and acetylation (Figure 5D). This provides a mechanistic link between HDAC6, Rictor, and FoxO1 regulation in the context of *P. gingivalis*-stimulated inflammatory responses (Figure 5D).

### Inhibition of FoxO1 Decreases NF-κB Activity and Inflammatory Cytokine Production in *P. gingivalis*-Stimulated BMDMs

Given that HDAC6 deficiency promotes the cytoplasmic retention of FoxO1 in *P. gingivalis*- stimulated macrophages, we next investigated whether FoxO1 is essential for NF-κB activation and the production of inflammatory mediators. As shown in Figure 6A, inhibition of FoxO1 using a specific inhibitor, AS1842856 (100 nM), largely reduced *P. gingivalis*-induced phosphorylation of NF-κB at Ser536 at various time points. Furthermore, pretreatment with AS1842856 significantly decreased the production of pro-inflammatory cytokines, including TNFα, IL-6, and IL-12p40, in *P. gingivalis*-stimulated BMDMs (Figures 6B). Interestingly, in Rictor-deficient mouse embryonic fibroblast (MEF) cells, depletion of Rictor led to a significant increase in the production of these inflammatory cytokines (Figures 6C-E). This observation supports the role of Rictor as a critical upstream regulator of FoxO1 activity in *P. gingivalis*-stimulated BMDMs. Altogether, these findings suggest that FoxO1 is essential for NF-κB activation and the subsequent production of inflammatory cytokines in response to *P. gingivalis*. Moreover, the regulation of FoxO1 activity by Rictor further highlights the interplay between upstream signaling pathways and FoxO1 in modulating inflammatory responses in macrophages.

**Figure 6.**
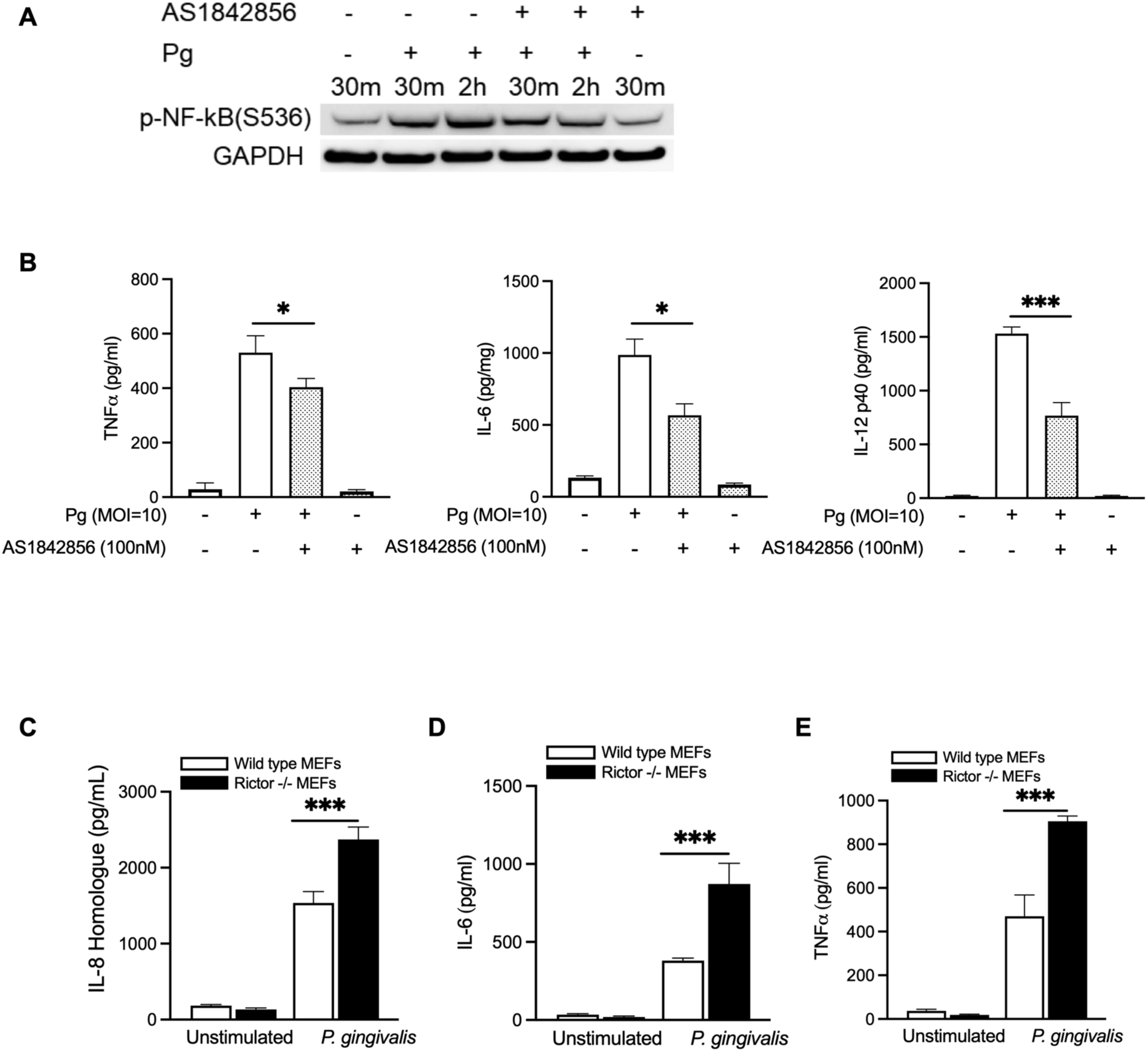
FoxO1 inhibition alleviates while Rictor depletion augments the production of pro-inflammatory cytokines in *P. gingivalis* stimulated immune cells. (A) BMDMs were pretreated with the FoxO1 inhibitor AS1842856 (100 nM) for 1 hour prior to *P. gingivalis* infection. Whole-cell lysates were collected at the indicated time points and analyzed by Western blot using the specified antibodies. **(B)** BMDMs were pretreated with AS1842856 for 2 hours, followed by an 8-hour *P. gingivalis* challenge. Cytokines (TNFα, IL-6 and IL-12P40) was measured by ELISA. **(C to E)** Rictor knockout (KO) MEFs were infected with *P. gingivalis* for 8 hours. IL-8 homologue (MIP2/CXCL2) **(C)**, IL-6**(D)** and TNFα **(E)** chemokine levels in the supernatants were quantified using ELISA. *, **, and ***, represent the statistical significance with *P* < 0.05, *P<0.01,* and *P < 0.001,* respectively.

### HDAC6 Deficiency Reduces Periodontal Inflammation and Alveolar Bone Loss in Mouse Periodontitis Models

Since we have demonstrated that HDAC6 depletion significantly attenuates inflammatory responses in cultured cells, we next evaluated its effects on periodontal inflammation and subsequent alveolar bone loss using ligature-induced and *P. gingivalis* infection-induced periodontitis models. For the ligature-induced periodontal bone loss model (Figure 7A), fourteen days after ligature placement, we examined the inflammatory status of gingival tissues and alveolar bone loss using micro-CT imaging system. As shown in figure 7B and C, HDAC6 KO mice were found to be significantly protected against ligature-induced bone loss in the presence and absence of *P. gingivalis* infection, as evidenced by a lower cementum-enamel junction to alveolar bone crest (CEJ-ABC) distance, as compared to wild-type controls (Figures 7B and 7C). These results are also confirmed by using the *P. gingivalis* infection model, showing HDAC6 KO significantly reduces oral inflammation-induced alveolar bone loss (Figure 7D and 7E). Moreover, the gingival tissues from the mice that were orally infected with *P. gingivalis* six times were also harvested 42 days post-infection for inflammation, following procedures described in previous studies (16, 41). As shown in Figures 7F, HDAC6 KO mice exhibited significantly reduced neutrophil infiltration in gingival tissues compared to wild-type controls. Immunohistochemical analysis (Figure 7G, H) revealed a reduction in nuclear FoxO1 accumulation and an increase in CD206 expression in HDAC6-deficient gingival tissues, indicating a shift toward an anti-inflammatory phenotype. These findings strongly suggest that HDAC6 deficiency reduces inflammation *in vivo*. Altogether, these results demonstrate that HDAC6 deficiency protects against inflammation-mediated alveolar bone loss *in vivo* by reducing periodontal inflammation and modulating the inflammatory response, highlighting its potential as a therapeutic target for periodontitis.

**Figure 7.**
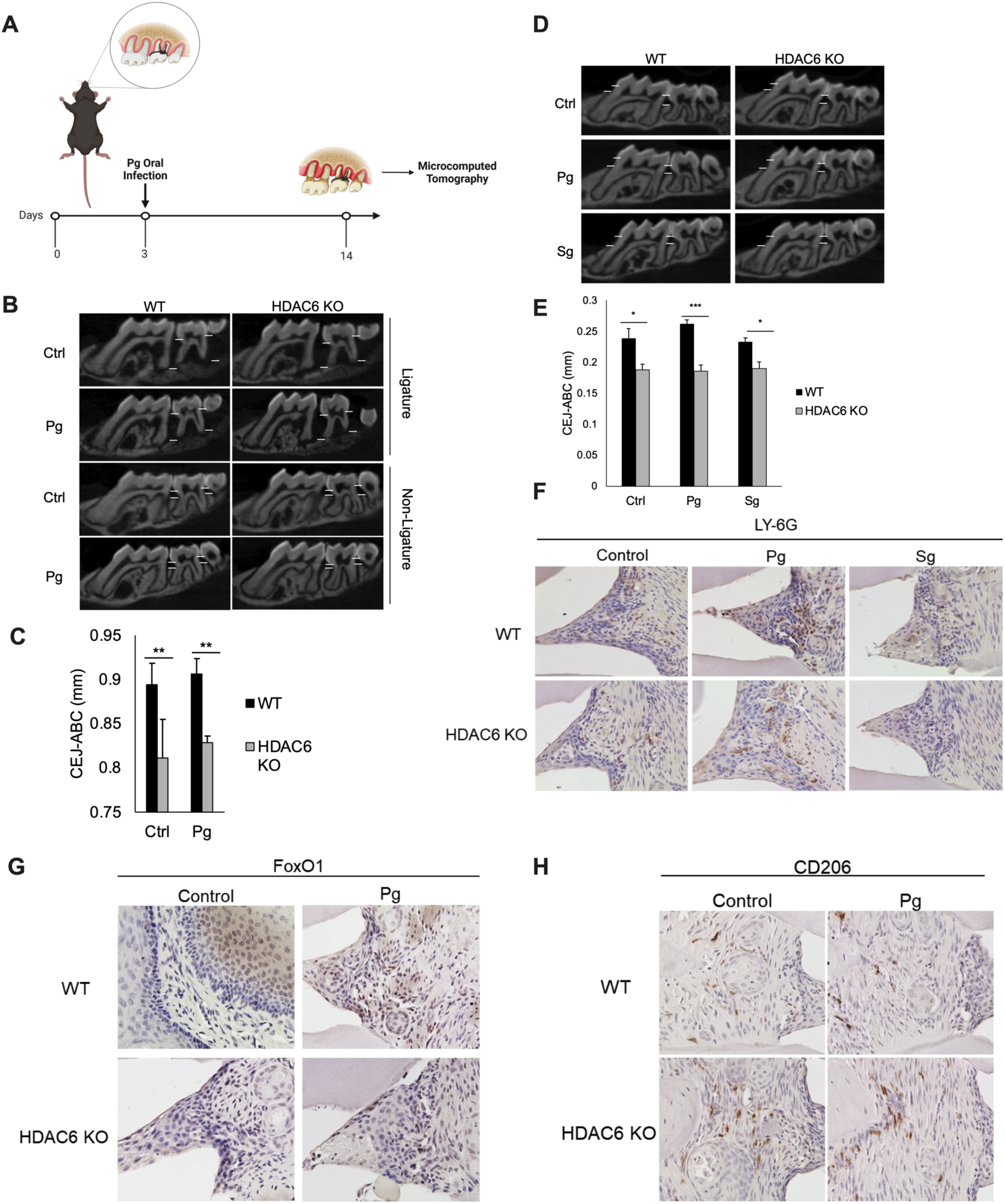
HDAC6 reduces periodontal inflammation and alveolar bone loss in mouse periodontitis models. (A-C) WT and HDAC6 KO mice (n = 5) were subjected to a ligation- induced periodontitis model. **(A)** Schematic illustration of the process for the ligation-induced periodontitis model. **(B)** Representative 2D images of alveolar bone loss, with measurements of the CEJ-ABC distance. **(C)** Quantification of bone loss via the CEJ-ABC distance on the left side of the M1 and M2 molars in ligation-induced periodontitis mice. **(D-H)** Oral infection-induced periodontitis model. **(D)** Representative 2D micro-CT images showing alveolar bone loss. Quantification of bone loss based on the CEJ-ABC distance for each side of the M1 and M2 molars. Results represent the average of palatal and buccal sides across all experimental mice in each group. Gingival tissues collected from mice (n = 5) orally infected with *P. gingivalis*, *S. gordonii*, or sham control were stained for LY-6G (**F**), FoxO1 (**G**), and CD206 (**H**) , and imaged using a Keyence microscope (60×).

## Discussion

Interaction and synergy between distinct post-translational modifications (PTMs), such as acetylation and phosphorylation, and their associated signaling networks remain poorly understood, particularly in the context of bacterial infection. In this study, utilizing periodontitis models, we demonstrate for the first time that HDAC6 is essential for regulating both the phosphorylation and acetylation of FoxO1, highlighting its critical role in *P. gingivalis*-induced periodontal inflammation. Furthermore, we found that *P. gingivalis* infection activates HDAC6, which promotes inflammatory responses through the concurrent modulation of FoxO1 acetylation and phosphorylation, unveiling a novel mechanistic module for bacterial infection-mediated inflammation (Figure 8).

**Figure 8.**
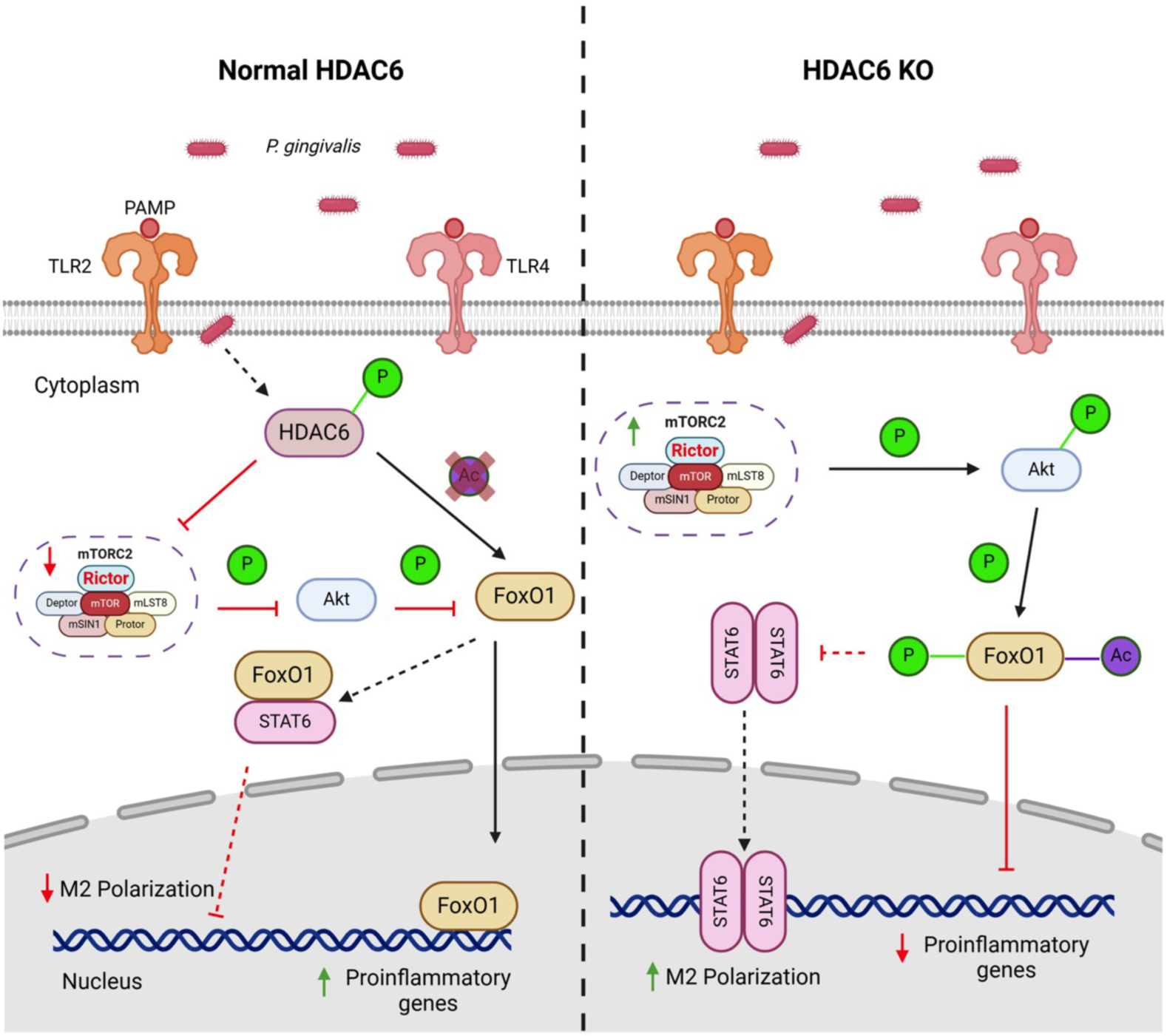
Proposed model of HDAC6 regulation of FoxO1 activity. In wild-type cells, *P. gingivalis* infection activates HDAC6 via phosphorylation. Activated HDAC6 then suppresses Rictor (mTORC2) degradation, thereby limiting the downstream phosphorylation of Akt and FoxO1, which facilitates FoxO1’s nuclear retention. Simultaneously, HDAC6 directly deacetylates FoxO1, further promoting its nuclear translocation. Through these two distinct mechanisms, the increased nuclear FoxO1 binds to DNA and induces the transcription of pro-inflammatory mediators. FoxO1 may also interact with STAT6, preventing its nuclear translocation and limiting M2 macrophage activation. In contrast, HDAC6 depletion enhances Rictor (mTORC2) expression and increases Akt and FoxO1 phosphorylation. Combined with increased FoxO1 acetylation, these concurrent modifications lead to its cytoplasmic sequestration, thereby inhibiting pro-inflammatory gene transcription. HDAC6 depletion also promotes STAT6 dimerization and DNA binding, facilitating M2 polarization. Red bar=inhibition; red arrow=downregulation; green arrow=upregulation; dotted arrow=possible activation; dotted bar=possible inhibition. Figure made in BioRender.

We observed that inhibition or deletion of HDAC6 increased FoxO1 acetylation and phosphorylation in response to *P. gingivalis* infection, enhancing its cytoplasmic sequestration and thereby suppressing the transcription of downstream pro-inflammatory cytokines. Notably, we identified that HDAC6 regulates FoxO1 phosphorylation by modulating the expression of Rictor, a key component of the mTORC2 complex, and subsequently Akt phosphorylation. However, the precise mechanisms by which *P. gingivalis* activates HDAC6 remain unknown. *P. gingivalis* is known to activate multiple Toll-like receptors (TLRs), including TLR2, TLR4, and TLR5, through virulence factors such as lipopolysaccharide (LPS), peptidoglycan, and flagellin (39). Additionally, *P. gingivalis* secretes gingipains, lysine- and arginine-specific proteases, that have been shown to activate protease-activated receptor 2 (PAR2). These factors likely contribute to the activation of HDAC6 during infection. HDAC6 activity is regulated by phosphorylation at specific amino acid residues, and our data show that phosphorylation enhances HDAC6 activation (Figure 2A, C; Figure 4B). Interestingly, while *P. gingivalis* LPS induces HDAC6 activation, it does so less robustly than whole bacteria, and *E. coli* LPS fails to activate HDAC6 entirely (Data now shown). These findings suggest that *P. gingivalis*-mediated activation of HDAC6 may depend on TLR2 rather than TLR4, as TLR2 is the primary receptor activated by *P. gingivalis*. This hypothesis aligns with previous studies identifying ASK1 kinase and Rac1 as critical mediators of HDAC6 activity (70, 71), both of which are reported to be activated upon TLR2 activation. TLR2 signaling through Rac1, PI3K, and Akt has been shown to activate NF-κB in macrophages and neutrophils (72, 73), suggesting that Rac1 may link TLR2 signaling to HDAC6 activation. Rac1 has previously been implicated in HDAC6-mediated functions (51, 71), such as regulating cell migration and invasion, indicating that Rac1 may act as a signaling node connecting TLR2 and HDAC6 signaling in response to *P. gingivalis*. Additionally, our findings further reveal that HDAC6 regulates Akt phosphorylation through the ubiquitination-dependent modulation of Rictor expression. Given that Rac1-mediated Akt phosphorylation is well-documented, it is highly possible that HDAC6 represents a secondary branch of Rac1 signaling, promoting Akt activation by modulating Rictor expression. Another plausible mechanism involves gingipain-mediated degradation of HDAC6 suppressors. For instance, the 14-3-3 protein, a known HDAC6 inhibitor, could be proteolyzed by secreted gingipains, thereby relieving constraints on HDAC6 activity. Supporting this idea, our unpublished data show that *P. gingivalis* mutants lacking lysine-specific gingipain (ΔKgp) fail to activate HDAC6 or induce FoxO1 deacetylation. Lastly, HDAC6 activity is regulated by ERK1, which binds and phosphorylates HDAC6 at specific residues, promoting its deacetylation of substrates like α-tubulin. ERK1 activation has been widely reported in response to *P. gingivalis* infection, either directly or indirectly via pro-inflammatory cytokines such as TNF- α and IL-6 (74, 75). While ERK1 activation is common to multiple bacterial infections, such as *E. coli*, only *P. gingivalis* effectively activates HDAC6, suggesting it is not a predominate mechanism in this context. Altogether, our findings underscore the need for further studies to dissect the components of *P. gingivalis* involved in HDAC6 activation and the signaling pathways involved. Such investigations will be crucial for unraveling the intricate regulatory networks modulating inflammatory responses during bacterial infection and may inform the development of novel therapeutic strategies for periodontal and other inflammatory diseases.

In this study, we found that HDAC6 depletion promotes macrophage polarization toward the anti-inflammatory M2 phenotype in response to *P. gingivalis* infection. Furthermore, HDAC6 deficiency suppresses FoxO1 nuclear retention and diminishes its transcriptional activity. Previous studies, including our own, have demonstrated that FoxO1 is involved in macrophage polarization (16, 17), suggesting that FoxO1 may influence this process by regulating the activity of other transcription factors (TFs). Using AlphaFold3 for protein-protein interaction modeling, our results indicate a high likelihood of direct binding between FoxO1 and STAT6 (Figure S1). The predicted template modeling (pTM) score of 0.77, which reflects high structural similarity, supports the validity of this prediction. This confidence is further reinforced by the comparable molecular weights of FoxO1 (∼110 kDa) and STAT6 (∼94 kDa), which reduce the potential for size-related biases in the prediction. As we know, STAT6 is a key transcription factor driving M2 macrophage polarization. Based on our findings, we propose that FoxO1 may bind STAT6 and suppress its activity through an as-yet-unknown mechanism. Consistent with this, FoxO1 has been shown to promote M1 polarization through direct binding to pro-inflammatory gene promoters, such as *il- 1β* (76, 77). Our modeling data provide the first evidence of a direct interaction between FoxO1 and STAT6, aligning with previous reports suggesting a suppressive effect of FoxO1 on STAT6 activity (78). Notably, STAT6 monomers are phosphorylated by Janus kinases (JAKs) in response to IL-4 or IL-13, enabling their dimerization and subsequent nuclear translocation to drive M2-associated gene transcription. In pro-inflammatory conditions, such as *P. gingivalis* infection, elevated FoxO1 levels may disrupt this process by competitively binding STAT6, preventing its phosphorylation, dimerization, or ultimately DNA binding. Additionally, nuclear FoxO1 may directly interact with STAT6 to inhibit its transcriptional activity. HDAC6 depletion, which reduces FoxO1 activity, likely frees STAT6 to efficiently dimerize, translocate to the nucleus, and drive M2-associated gene expression. Future studies employing LC-MS/MS to detect direct interactions between FoxO1 and STAT6, protein crystallography to observe their binding in solution, and chromatin immunoprecipitation (ChIP) assays to evaluate STAT6 DNA binding in WT and FoxO1-deficient macrophages will validate this structural prediction.

While we observed that HDAC6 depletion reduces periodontal inflammation and protects against alveolar bone loss in two distinct animal models, the effect of *P. gingivalis* infection on bone loss was less pronounced compared to non-infected controls. Periodontal inflammation is considered a primary driver of alveolar bone loss, yet *P. gingivalis* alone is a relatively weak inducer of inflammation *in vitro*. Its significance as a model organism stems from its ability to disrupt microbial homeostasis, leading to dysbiosis and amplifying the pathogenic potential of the entire oral microbiota(31, 32, 79). In our study, *P. gingivalis* infection was employed only as an inflammation trigger in both oral gavage and ligature models. In this regard, for the ligature model, the mechanical irritation caused by the ligature induces robust inflammation, potentially overshadowing the contribution of *P. gingivalis*-induced inflammation. Additionally, *P. gingivalis* alters the oral microbiome, potentially promoting colonization by bacteria with opposing effects on bone metabolism, which could offset the direct impact of *P. gingivalis* on alveolar bone loss through incrementing some inflammation. Furthermore, *P. gingivalis* induces both pro- and anti-inflammatory responses, such as activation of JAK3 and SGK1, which may temper localized inflammation and mitigate bone loss. In the oral gavage model, inflammation is chronic and relatively low-grade due to the extended experimental duration (>6 weeks). Factors such as infection frequency, cytokine dynamics, and the timing of sample collection may influence the observed phenotype. Therefore, in this study, *P. gingivalis* primarily served to amplify inflammation in WT and HDAC6 KO mice, allowing us to evaluate the role of HDAC6. Consistent with our hypothesis, *P. gingivalis* infection exacerbated inflammation in WT mice, whereas HDAC6 deficiency significantly reduced inflammation and inflammation-induced bone loss (Figure 7F). These findings, combined with results from the ligature model, underscore the critical role of HDAC6 in mediating periodontal inflammation and its downstream effects on alveolar bone loss.

In summary, this study is the first to reveal the pivotal role of HDAC6 in regulating periodontal inflammatory responses. We demonstrated that HDAC6 depletion suppresses the production of pro-inflammatory cytokines, shifts macrophage polarization toward anti-inflammatory M2 phenotypes, and protects against *P. gingivalis*-induced alveolar bone loss. Furthermore, our findings highlight a dual mechanism by which HDAC6 modulates inflammation: through the simultaneous regulation of FoxO1 acetylation and phosphorylation, promoting its cytoplasmic retention and attenuating its transcriptional activity. This two-pronged strategy provides novel insights into the molecular pathways governing periodontal inflammation. Given the recent FDA approval of HDAC inhibitors for treating various diseases and the ongoing development of selective inhibitors targeting specific HDAC isoforms, future efforts to further investigate HDAC6-specific inhibitors could lead to effective treatments not only for periodontal disease but also for other inflammatory conditions where HDAC6 and FoxO1 play central regulatory roles.

## Acknowledgments

This research was supported by Grants R01 DE026727 (H.W.), R21 DE031376 (H.W.), and F31 DE031968 (to H. L.) from the U.S. National Institute of Dental and Craniofacial Research, VCU Breakthroughs and Presidential Research Quest Funds (PeRQ) award to HW.

## Authors’ Contribution

H. Lohner, X. Han, J. Ren, R. Liang and H. Wang performed research; H. Lohner and H. Wang analyzed the data; R. Liang performed the Alpha3 protein binding predictions; H. Wang, R. Liang, and S. Liang designed and supervised the research; H. Wang wrote the draft of the manuscript; H. Lohner and S. Liang reviewed and proofread the manuscript. All authors orally approved to deposit this preprint version to the bioRxiv.

## Disclosure

All authors have reviewed and approved this manuscript for deposition as a preprint on bioRxiv.

This manuscript has not been accepted or published elsewhere.

The authors declare no competing interests.

## Supplemental figures

**Fig. S1.**
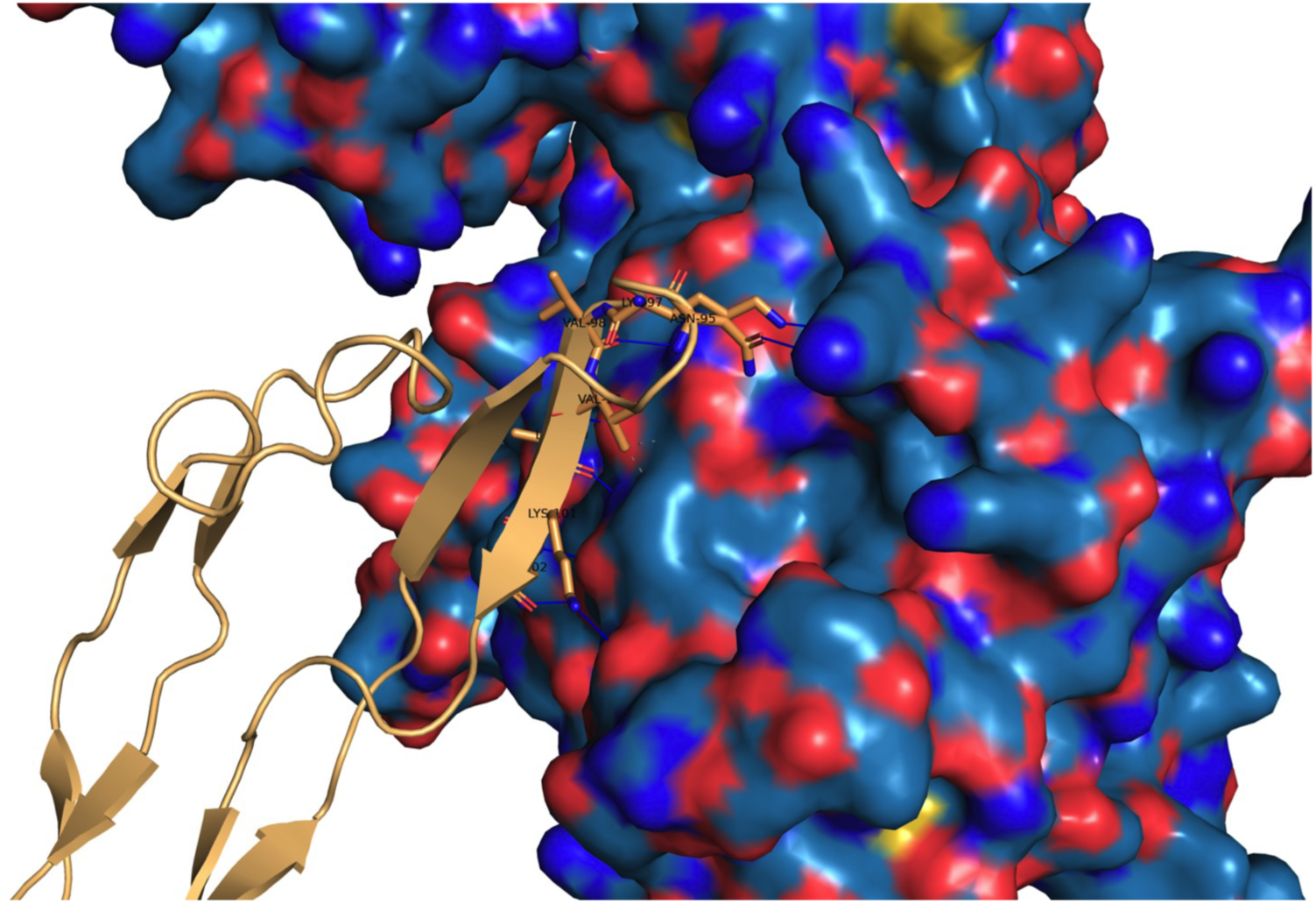
AlphaFold 3 prediction of FoxO1-STAT6 complex. Surface, STAT6 Ser2-Gln654; bronze ribbon and sticks, FoxO1 Lys151-Ser266. Blue line, hydrogen bond; yellow dash, salt bridge; grey dash, hydrophobic interaction. Residue labels are FoxO1 counting from Lys1 to Ser116. There are 28 hydrogen bonds, 2 salt bridges, and 1 hydrophobic interaction between FoxO1 and STAT6. The structure is rendered with PyMOL. Interactions were analyzed with protein-ligand interaction profiler. Interface predicted template modeling score (ipTM)=0.23; predicted template modelling score (pTM)=0.77.

